# High confidence identification of intra-host single nucleotide variants for person-to-person influenza transmission tracking in congregate settings

**DOI:** 10.1101/2021.07.01.450528

**Authors:** Irina Maljkovic Berry, Todd Treangen, Christian Fung, Sheldon Tai, Simon Pollett, Filbert Hong, Tao Li, Papa Pireku, Ashley Thomanna, Jennifer German, P. Jacob Bueno de Mesquita, Wiriya Rutvisuttinunt, Michael Panciera, Grace Lidl, Matthew Frieman, Richard G Jarman, Donald K Milton, Prometheus@UMD Consortium

**Affiliations:** Walter Reed Army Institute of Research, Silver Spring, MD, USA; Rice University, Department of Computer Science, Houston, TX, USA; University of Maryland School of Public Health, Institute of Applied Environmental Health, College Park, MD, USA; University of Maryland School of Medicine, Department of Microbiology and Immunology, Baltimore, MD, USA

## Abstract

Influenza within-host viral populations are the source of all global influenza diversity and play an important role in driving the evolution and escape of the influenza virus from human immune responses, antiviral treatment, and vaccines, and have been used in precision tracking of influenza transmission chains. Next Generation Sequencing (NGS) has greatly improved our ability to study these populations, however, major challenges remain, such as accurate identification of intra-host single nucleotide variants (iSNVs) that represent within-host viral diversity of influenza virus. In order to investigate the sources and the frequency of called iSNVs in influenza samples, we used a set of longitudinal influenza patient samples collected within a University of Maryland (UMD) cohort of college students in a living learning community. Our results indicate that technical replicates aid in removal of random RT-PCR, PCR, and platform sequencing errors, while the use of clonal plasmids for removal of systematic errors is more important in samples of low RNA abundance. We show that the choice of reference for read mapping affects the frequency of called iSNVs, with the sample self-reference resulting in the lowest amount of iSNV noise. The importance of variant caller choice is also highlighted in our study, as we observe differential sensitivity of variant callers to the mapping reference choice, as well as the poor overlap of their called iSNVs. Based on this, we develop an approach for identification of highly probable iSNVs by removal of sequencing and bioinformatics algorithm-associated errors, which we implement in phylogenetic analyses of the UMD samples for a greater resolution of transmission links. In addition to identifying closely related transmission connections supported by the presence of highly confident shared iSNVs between patients, our results also indicate that the rate of minor variant turnover within a host may be a limiting factor for utilization of iSNVs to determine patient epidemiological links.

## INTRODUCTION

Influenza pandemics, epidemics and outbreaks have caused significant morbidity and mortality in the human population. Despite the availability of influenza vaccines, the virus infects an approximate 10% of the global population annually, causing 290 000–650 000 deaths each year (1, 2). The persistence and the epidemiological success of influenza virus is mainly attributed to its rapid evolution, through which the virus continuously attains mutations resulting in escape from the immune system, vaccines, and antivirals. However, this rapid evolution has also enabled numerous genomic studies resulting in critical insights into the dynamics of influenza evolution, transmission and spread (3–10).

The high evolutionary rate of the influenza virus is driven by selection pressures imposed on the virus during its replication cycle within a host. Because of its error-prone RNA polymerase, the majority of the newly produced viral genomes will differ by one or more mutations, creating an underlying population of viral variants from which, at any time point, the fittest ones can be selected for further propagation. The frequency of viral variants within an intra-host viral population is determined not only by selection, but also by evolutionary bottlenecks (such as transmission and population immunity), and stochastic events (11). Therefore, within a host, influenza exists not as a single virus, but as a dynamic population of variants providing plasticity to rapidly adapt to environmental changes.

The importance of within-host viral populations and their variants has been demonstrated for numerous rapidly evolving viruses, including influenza. These intra-host viral populations are the very source from which all viral global diversity stems, and are also the source from which all viral escape variants are selected. For instance, presence of drug resistant minor variants within a host has been associated with treatment failure, and the mechanisms of such emergence have been studied, leading to development of methods for measurement of resistance based on frequencies of intra-host viral variants (12–17). Furthermore, transmission of drug-resistant intra-host minor variants has been observed in influenza, resulting in their rapid increase and fixation within the recipient host following treatment (16). Intra-host viral dynamics and diversity have also been associated with disease progression and disease severity (18–20). Importantly, within-host variants have been associated with escape from human immune system as well as escape from vaccine-induced immune responses, leading to development of methods that take into account intra-host viral fitness to inform design of immunogens and therapies (21–25). The presence and frequency of intra-host single nucleotide variants (iSNVs) has also been used to investigate between-host transmission events and refine phylogenetic clustering (6, 26–32). In this case, the iSNVs provide greater resolution of transmission events, as consensus whole genome sequences typically have insufficient variability to distinguish between infecting strains sampled within two weeks of each other, especially if closely sampled in space (33–35).

The ubiquity of Next Generation Sequencing (NGS), with its ease of whole genome sequencing and its high sequencing depths, has greatly expanded our ability to identify and study intra-host viral populations and their variants. However, despite the recent advances in the field, accurate characterization of iSNVs and intra-host variant populations still remains a challenge. Many sources of error exist that contribute to the calling of false positive iSNVs within a dataset, such as iSNVs generated due to the errors in RT-PCR and PCR, inherent sequencing errors of different sequencing platforms, base calling errors, primer-induced errors, and biases introduced by genome assembly and basecalling algorithms (36). Approaches such as the use of unique molecular identifiers, circular sequencing, and duplex sequencing, have been developed to mitigate some of the sequencing-associated errors, and to provide more accurate iSNV identification (37). More recently, sequencing of sample technical replicates and low-variant controls has been deployed for intra-host population analyses of SARS-CoV-2 (38, 39). However, to date, no study has comprehensively investigated iSNV errors introduced by all the above approaches, and more importantly, no study has accounted for all of the errors in their iSNV identification and analysis. In this study, we use a sample set from influenza surveillance at University of Maryland dorms to investigate the magnitude and the characteristics of different types of iSNV errors, to remove false positive variants, and to reconstruct fine scale virus transmissions within UMD dorms by utilizing high-confidence iSNVs. This framework for transmission tracking has major public health relevance, for instance, for implementation of the reconstructions of influenza and other respiratory virus spread in congregate settings such as college dormitories, military training settings, and ships.

## RESULTS

Eighteen influenza A and B samples (Table 1), obtained from the season 2017-2018 respiratory surveillance at the University of Maryland, as well as three contemporary influenza A and B plasmids, were sequenced and analyzed as described in Materials and Methods (Table 1, Figure S1). All the sequenced and assembled genomes and positions with called iSNVs at 1% or higher had a depth of read coverage >1000x and all segments were sequenced for each of the samples.

**Table 1.**
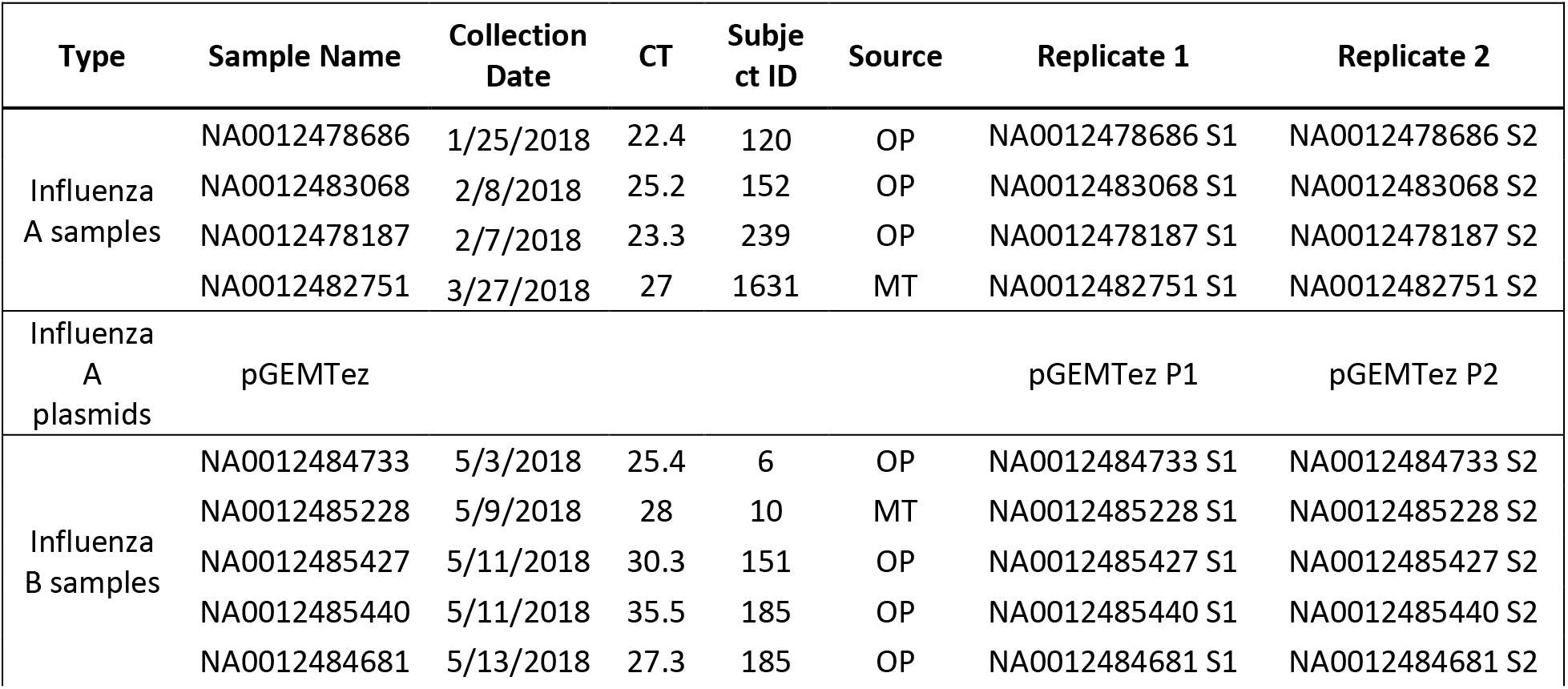
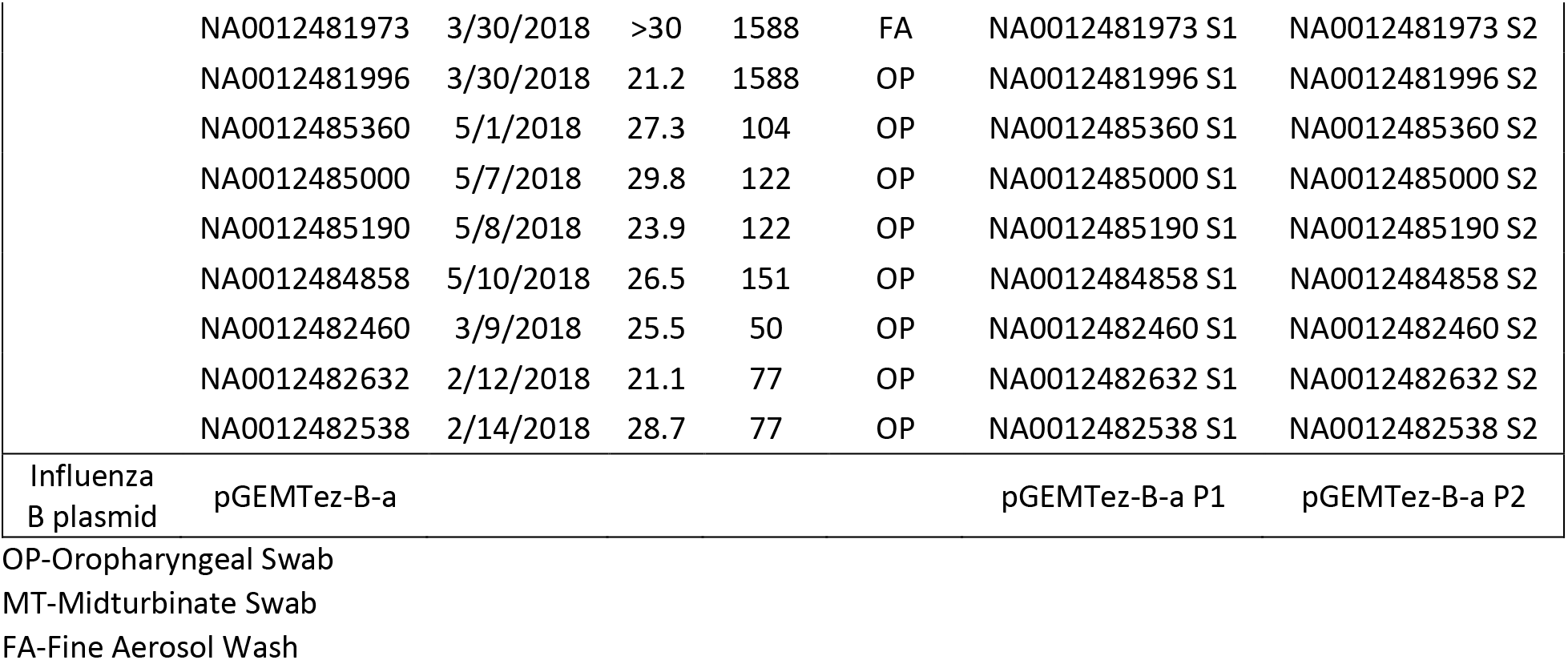
Sequencing of samples and plasmids.

Two technical replicates (S1 and S2) of each sample were used to confirm iSNVs and remove random RT-PCR, PCR and sequencing errors. Inclusion of plasmid (P) was to account for any systematic errors during the sequencing process. To test the importance of the reference choice for iSNV calling, three different reference whole genome sequences with increasing genetic distance from the sample, self-reference (construction of sample consensus by initial mapping to a lineage/subtype reference, followed by a re-run with read mapping to the own consensus genome), B/Victoria (CY249062.1-CY249069.1), and B/Yamagata (MH584285-MH584292), were used for reference-guided genome assembly. Finally, three different variant callers (ngs_mapper, LoFreq and iVar) were used for variant calling to assess variant caller-induced error. The called iSNV positions were plotted: i) for each sample replicate separately (S1, S2), ii) for positions overlapping between the two replicates (S1+S2), iii) for each sample replicate separately excluding iSNV positions found in the plasmids (S1-P, S2-P), iv) for positions overlapping between the two sample replicates excluding positions found in the plasmids ((S1+S2)-P).

### Self-reference mapping with PCR/sequencing error removal is most reliable for iSNV calling

In order to investigate iSNVs called by the S1, S2, S1+S2, S1-P, S2-P and (S1+S2)-P approaches, and the impact of reference choice on these iSNV calls, the initial analyses of the raw (non-curated) data were performed on a single influenza sample, NA0012484733. The virus from this sample belonged to the Victoria lineage of influenza B.

Our results showed decreasing number of called iSNV positions when counting the positions confirmed by both sample replicates (S1+S2) compared to S1 and S2 alone, and also when taking into account plasmid information (S1-P, S2-P and (S1+S2)-P), indicating that these called iSNVs may be false positives, present in the sample due to random or systematic PCR and sequencing errors. In addition, our results showed that the number of called iSNV positions was sensitive to the reference strain used for read mapping, with the more divergent reference (B/Yamagata) resulting in most called iSNV positions, and the least divergent reference (self-reference) resulting in the lowest number of called iSNVs (Figure 1). Furthermore, comparisons between the ngs_mapper, LoFreq and iVar variant callers indicated that this reference-based sensitivity also depended on the algorithm of choice, with LoFreq being the most sensitive to reference divergence from the sample. Indeed, while LoFreq Yamagata mapping resulted in >800 iSNV positions for the sample NA0012484733 S1 and S2 replicates, which was the highest number of iSNV positions found among all variant callers, its self-reference mapping gave the smallest number of iSNVs among all three variant callers (Figure 1). These results indicated that using self-reference reduces most of the iSNV noise, and that using replicate sample sequencing for removal of random PCR and sequencing error and inclusion of plasmids for removal of systematic error ((S1+S2)-P) was important for calling of high confidence iSNVs in a sample, especially when more divergent reference was used for mapping.

**Figure 1.**
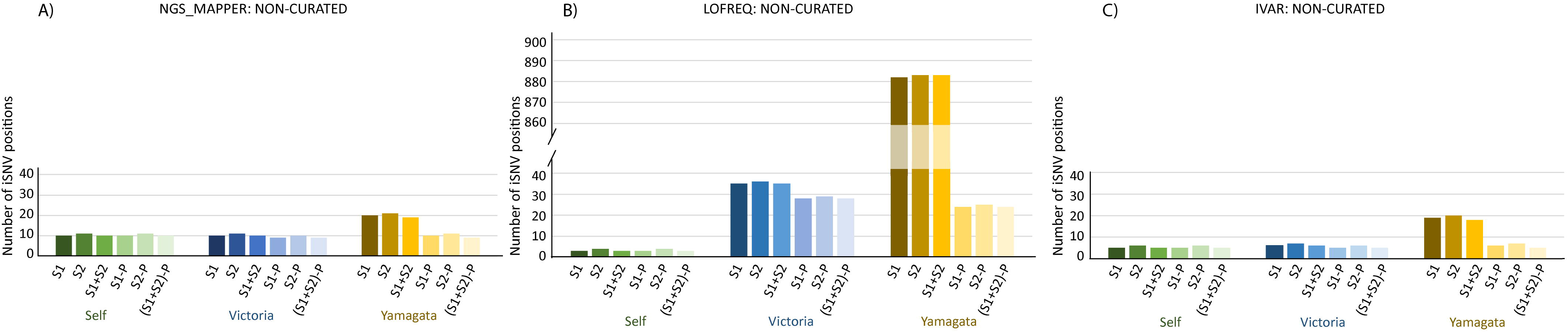
Number of iSNV positions in non-curated sample NA0012484733 when comparing sequencing, reference and algorithm-based errors. A) ngs_mapper variant caller; B) LoFreq variant caller; C) iVar variant caller. Green bars = self-reference; blue bars = B/Victoria reference; yellow bars = B/Yamagata reference. S1 = sample replicate 1; S2 = sample replicate 2; S1+S2 = iSNVs confirmed by both S1 and S2; S1-P = sample replicate 1 excluding iSNVs found in plasmids; S2-P = sample replicate 2 excluding iSNVs found in plasmids; (S1+S2)-P = iSNVs confirmed by both S1 and S2 excluding iSNVs found in plasmids.

### Variant caller-related errors contribute to calling of false iSNVs

Based on the above results, we compared the exact (S1+S2)-P iSNV positions called by the three different variant callers and references, to assess their called iSNV position overlap. This approach would remove all the potential sequencing and PCR errors, thus only highlighting variant calling algorithm errors. Our results showed great disparity between the variant callers and references (Figure 2). Thus, even when approximately the same numbers of iSNV positions were called, these did not overlap well between the variant calling algorithms, or within the same algorithm with divergent references. For sample NA0012484733, only three iSNV CDS positions were found (segment 1: 1868, segment 2: 1136 and segment 8: 821) that were confirmed by all the three variant callers and all three reference types. Interestingly, LoFreq with self-reference found only the three confirmed positions and no additional ones. In addition, when only comparing two algorithms to each other (ngs_mapper and LoFreq, ngs_mapper and iVar, or LoFreq and iVar) only the same three confirmed positions were found. These results indicate that these three positions were most probably true iSNV positions, confirmed by both duplicate samples, plasmids, all the references, and all the variant callers. Thus, by using a self-reference for mapping, utilizing information from at least two different variant callers, and removing the various sequencing errors ((S1+S2)-P), many false-positive iSNVs can be eliminated and iSNVs of high probability can be identified.

**Figure 2.**
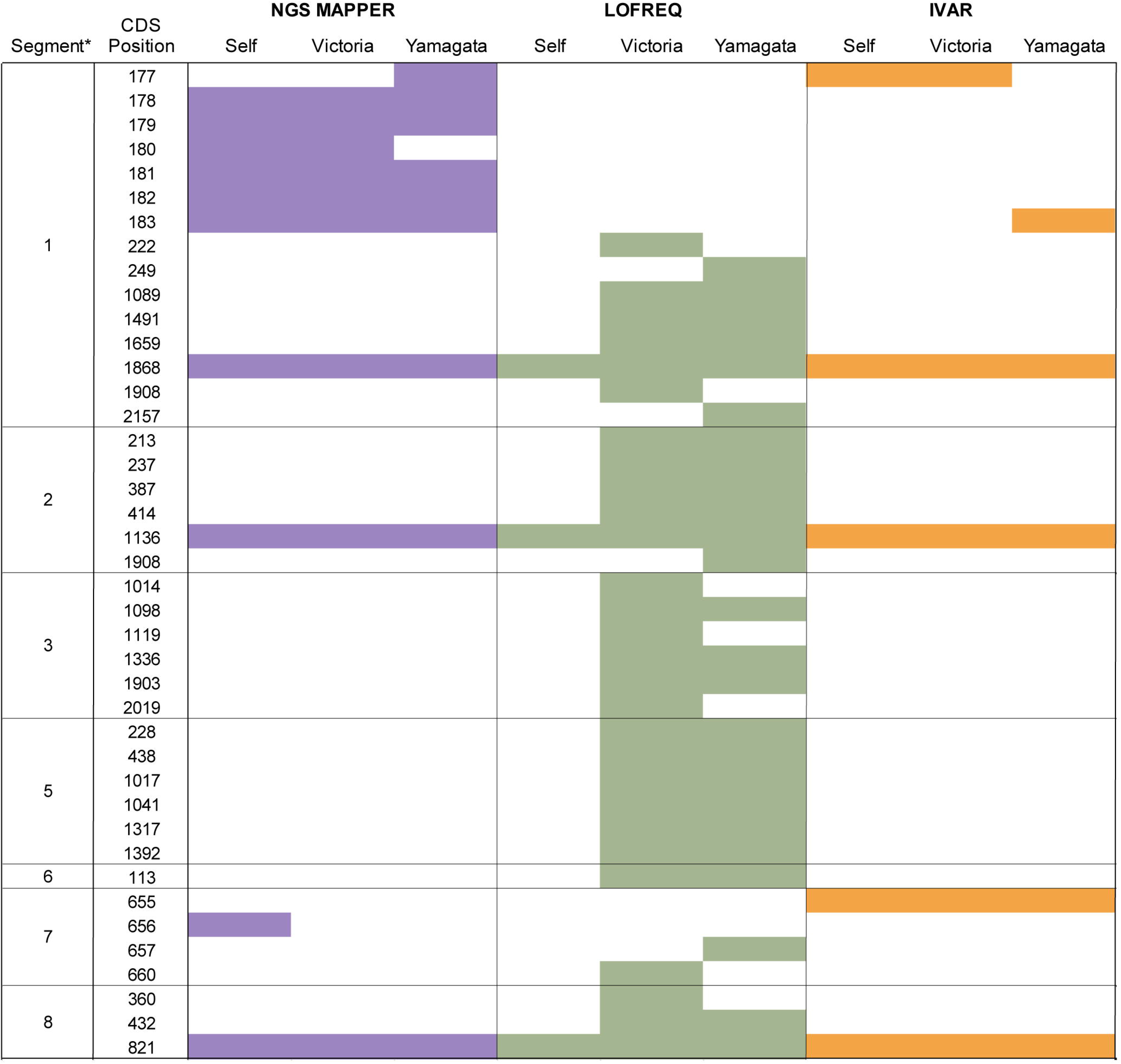
Sample NA0012484733 (S1+S2)-P iSNV positions. ngs_mapper (purple), LoFreq (green), and iVar (orange), and three references for read mapping. *Segment 4 did not have any iSNV positions.

In order to confirm the results from the single sample NA0012484733 analyses, we repeated the comparisons of iSNV positions found by different variant calling algorithms and sequencing error-reducing approaches on the complete set of the influenza A and B virus positive samples. We did this comparison using self-reference mapping only, as this was consistently found to produce the fewest number of called iSNV positions across the various NA0012484733 analyses (Figure 1) and to remove the most of the false positive noise signal. For the four influenza A virus samples (Figure 3A), and eight CT<30 influenza B virus Victoria lineage samples (Figure 3B), we observed the same patterns as in the initial single-sample self-reference results (Figure 1): using duplicate samples to remove sequencing and PCR errors did prove useful for removing false signal, while using plasmid information to remove systematic error was less consequential when using self-reference for read assembly. However, in the three influenza B Victoria lineage samples with CT>30, the observed patterns differed (Figure 3C). Here, using both duplicate samples and plasmid information was crucial for removal of various sequencing and PCR errors. The number of plasmid positions that contributed to the removal of error in the samples was significantly higher for influenza B Victoria CT>30 samples (28-36 positions) than in influenza B Victoria CT<30 samples (0-2 positions), p=1.96E-08, F-test. Similar patterns were also observed for the three influenza B Yamagata lineage samples (Figure 3D).

**Figure 3.**
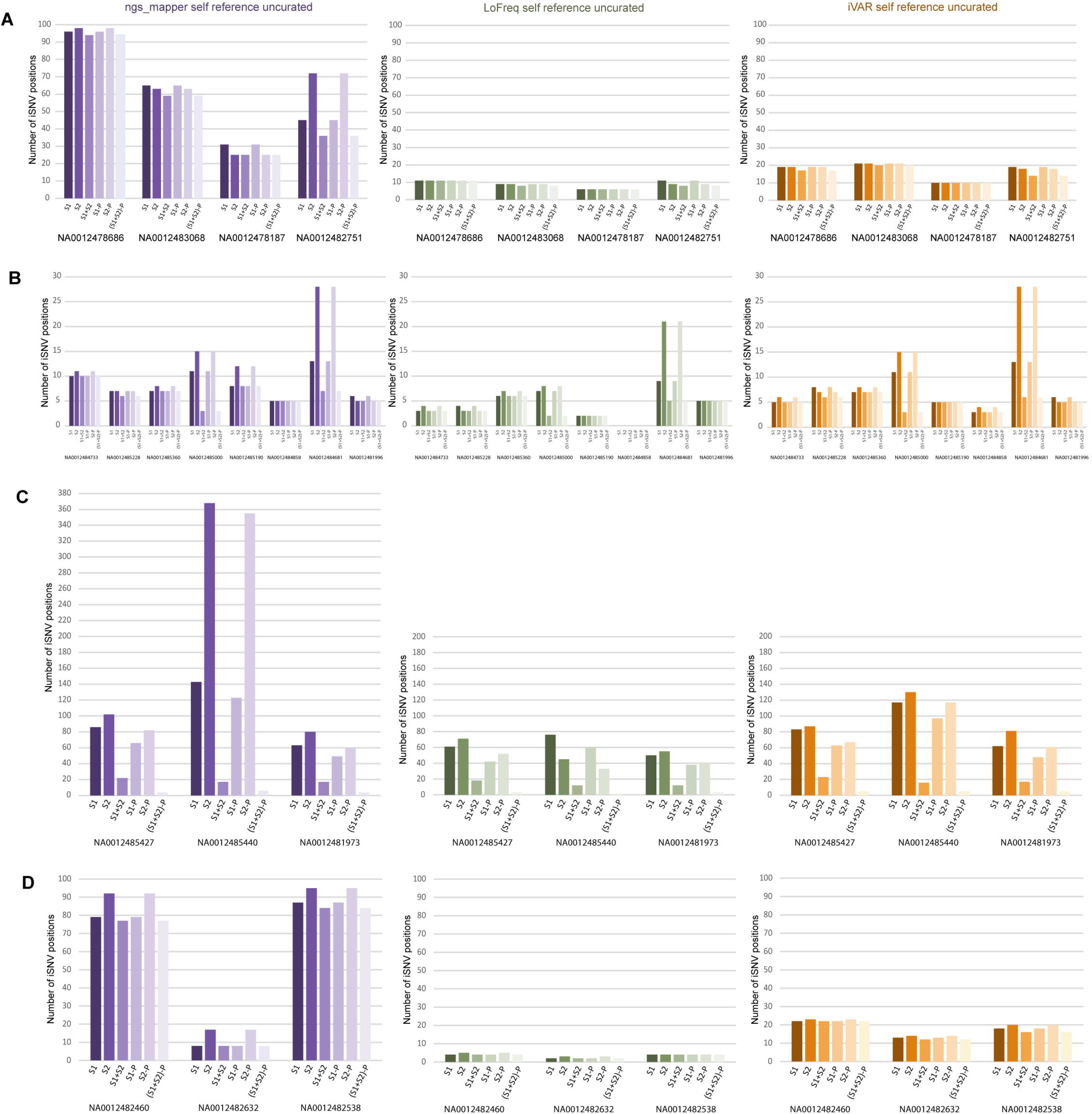
Number of iSNV positions in non-curated samples. A) influenza A, B) influenza B Victoria CT<30, C) influenza B Victoria CT>30, D) Influenza B Yamagata. Purple bars: ngs_mapper, green bars: LoFreq, orange bars: iVar.

The number of (S1+S2)-P positions was highest using ngs_mapper and lowest using LoFreq. Consistent with the results for the single sample, LoFreq found only positions that were also confirmed by both other variant callers, except for in one low viremia sample (Figure S2). These results show that LoFreq displays lowest amount of algorithm-associated error when using self-reference for mapping and removing sequencing and PCR errors. Confirming (S1+S2)-P positions by LoFreq and at least one other variant calling algorithm resulted in higher-confidence iSNV positions.

### Manual curation and consistency of iSNV frequencies

In order to assess the importance of manual genome curation and quality check, which takes into account strand bias and removes primer-induced errors, we compared the above uncurated results to the results from their manually curated data. We noted that manual data curation removed some iSNV positions from all the three variant callers, however, we note that the fewest iSNV positions were removed from LoFreq, which also had the smallest number of called positions to start with. This was consistent across different references (Figure S3) and samples (Figure S4). Therefore, to call high-confidence iSNV positions in our samples, we removed sequencing and PCR errors by duplicate sample and plasmid sequencing ((S1+S2)-P), we used self-reference for genome assembly, we removed algorithm-associated error by double-algorithm iSNV confirmation, and we removed primer –associated error by manual iSNV curation. The remaining iSNV positions were considered high-confidence, and were further analyzed for variant frequency comparisons. Our results indicated that variant frequencies in the highly confident iSNV positions were very consistent across algorithms and sample duplicates (Table 2 and Tables S1, S2). We also noticed that influenza B samples with low RNA abundance had fewer confirmed iSNV positions (average 0.33, range 0-1) than high RNA abundance samples (average 2.1, range 0-5).

**Table 2.** iSNV frequencies of sample NA0012484733 for three variant callers using the (S1+S2)-P approach and self-reference for mapping.

| Segment | CDS Position | LoFreq |  | ngs_mapper |  | iVar |  |
| --- | --- | --- | --- | --- | --- | --- | --- |
|  |  | S1 nt[aa] | S2 nt[aa] | S1 nt[aa] | S2 nt[aa] | S1 nt[aa] | S2 nt[aa] |
| 1 | 1868 | C[S]=42.36%<br>T[L]=57.64% | C[S]=42.55%<br>T[L]=57.45% | C[S]=51%<br>T[L]=49% | C[S]=51%<br>T[L]=49% | C[ ]=51.18%<br>T[L]=48.80% | C[ ]=50.71%<br>T[L]=49.26% |
| 2 | 1136 | A[N]=1.62%<br>G[S]=98.38% | A[N]=1.34%<br>G[S]=98.66% | A[N]=2%<br>G[S]=98% | A[N]=2%<br>G[S]=98% | A[N]=1.84%<br>G[S]=98.16% | A[N]=1.51%<br>G[S]=98.49% |
| 8 | 821 | A[Q]=1.11%<br>G[R]=88.29% | A[Q]=1.18%<br>G[R]=88.62% | A[Q]=1%<br>G[R]=99% | A[Q]=1%<br>G[R]=99% | A[Q]=1.25%<br>G[R]=98.75% | A[Q]=1.31%<br>G[R]=98.69% |

nt-nucleotide
aa-amino acid

### Influenza A and B virus transmission tracking through consensus and iSNV information

Consensus genome phylogenetic analyses of influenza A HA segments revealed at least three separate introductions of H3N2 into the study cohort during the 2017-2018 season (Figure 4A). Two of the samples clustered together in the tree, and also shared one iSNV in PB1 position 720, highlighting their close relationship and possible direct transmission of the virus between the individuals. For influenza B, 11 samples belonged to the Victoria lineage and were found in two well defined separate clusters, indicating at least two separate introductions of this virus into the cohort during the 2017-2018 season (Figure 4B). However, due to the high similarity of HA segments, the tree was mostly unresolved with low branch confidence values, and the number of introductions might have been higher. Even so, we found seven samples in a monophyletic cluster with high node support, showing their common ancestry and suggesting that the introduction of the virus was followed by onward spread within the study population. All seven samples were unresolved within this cluster, meaning that their true genomic relationships to each other could not be determined. An additional two samples were unresolved on the trunk of the tree just outside of this cluster, indicating that their relationship to the cluster sequences could not be resolved. However, we found that three of the samples within the monophyletic cluster shared iSNVs in PB1 position 1875, and two these samples also shared iSNV in NA position 509. The two samples that shared both PB1 and NA iSNVs were derived from the same participant (participant 122), sampled on two consecutive days. The third sample sharing an iSNV was from a different participant (participant 6), indicating close relationship of the viruses between these two individuals and possible direct transmission between the two. Three of the influenza B viruses from UMD samples belonged to the B/Yamagata lineage. Two were consecutive samples from the same participant and clustered together in the tree, and one was from a different participant clustering separately, indicating at least two separate introductions of influenza B Yamagata lineage into the UMD dorms (Figure S5).

**Figure 4.**
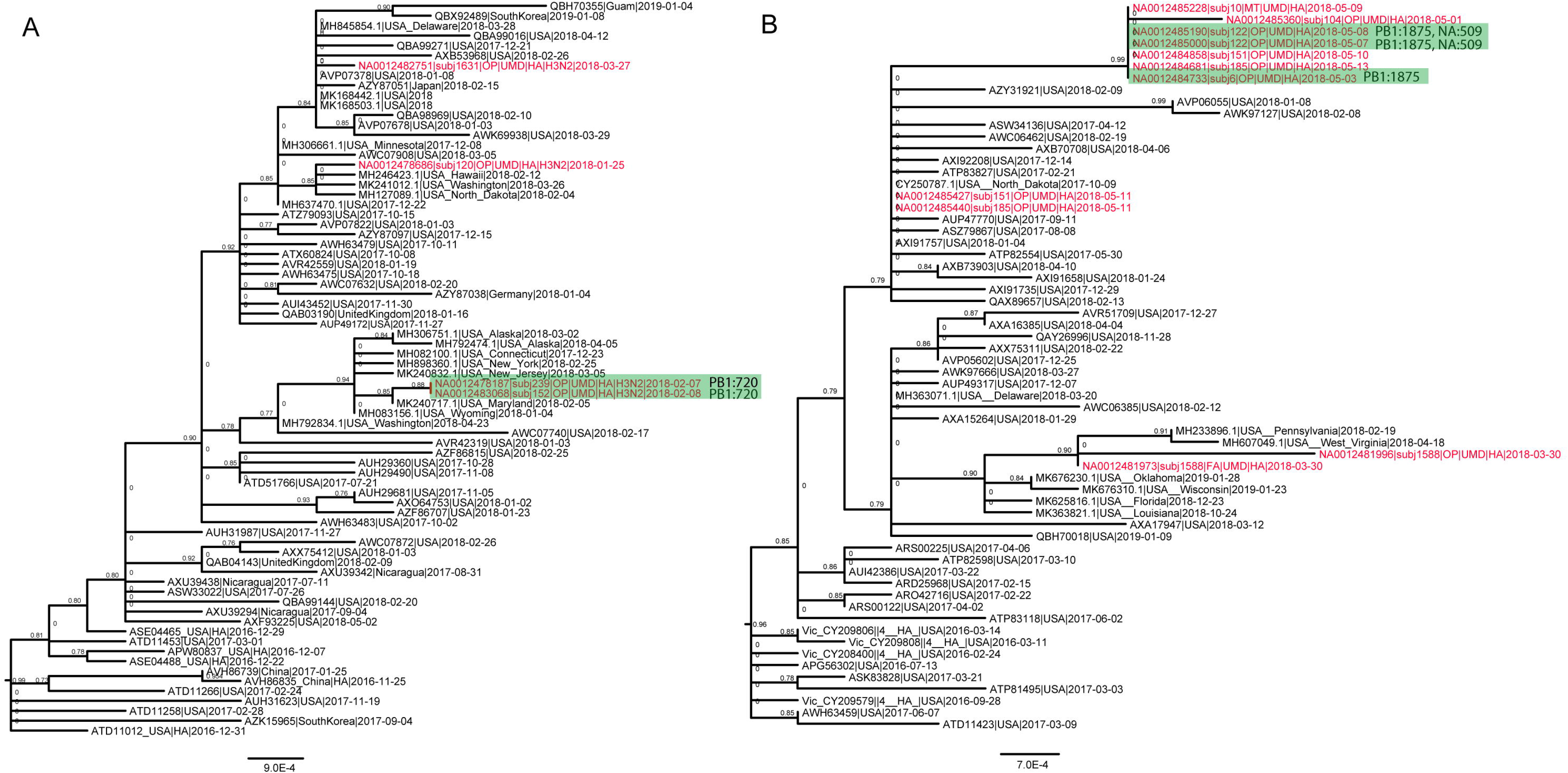
Maximum likelihood phylogenetic trees of the HA segment. A) Influenza A, B) Influenza B Victoria lineage. UMD samples are marked in red text and samples sharing iSNVs are marked with green boxes. Scale indicates nucleotide substitutions/site.

## DISCUSSION

The evolution of within-host viral populations, including composition and dynamics of their iSNVs, plays a fundamental role in many aspects of virus research, such as transmission, immune responses, and development of antiviral treatment and vaccines. However, iSNVs have notoriously been difficult to identify with high accuracy, due to many different sources of error introduction during RT-PCR and PCR amplification, sequencing, as well as genome assembly and variant calling.

Here, we investigate the sources and the magnitude of iSNV error, and we develop an approach for characterization of highly confident iSNV calls. We show that using sample technical replicates reduces the number of called iSNV positions and aids in removal of random PCR and sequencing errors, while using a clonal sample to remove any systematic errors is more important when low viral load samples are sequenced. The effects of PCR and sequencing errors are very well known, and numerous approaches to account for these errors have been used to call minor variants within a sample, such as using barcodes, technical replicates, and synthetic controls (37–41). The importance of synthetic controls or plasmids for removal of false positive iSNVs becomes especially important for studies using minor variants to track transmissions at a fine spatial scale, as these errors can be mistaken for shared variants between individuals, and thus erroneously suggest epidemiological links.

Our results further highlight the errors associated with bioinformatics algorithms and genome assembly where false positive minor variants can arise. It is well known that a closely related genome should be used for reference mapping-based genome assembly, however, sometimes it is hard to predict how related a reference is before the assembly. Here we show that using a sample self-reference results in the fewest called iSNV positions, and that the number of iSNV positions rises with the increased sample distance from the reference used. This result is not surprising, as read mapping to self-reference will increase the number of mapped reads that are most similar to the sample itself, and thus decrease the proportion of the reads with the minor variant. Some of these minor variants that fall below a certain threshold might be real, however, they may represent viral variants less fit for replication, or variants selected against by the human immune system.

Our results also indicate that various variant callers differ in their sensitivity to reference relatedness to the sample, which is something that needs to be taken into account for variant caller selection. More importantly, our results show that variant callers differ greatly in which iSNV positions they call, often disagreeing with each other. The variant caller that found positions confirmed by the other variant callers, and those positions only, was LoFreq with self-reference mapping. However, in rare instances LoFreq would also call an iSNV not confirmed elsewhere. Therefore, to remove all possible algorithm-based errors, a self-reference is preferred for read mapping, along with confirmation of minor variants with at least two different variant callers. Although this approach might result in removal of some true iSNV positions (false negatives), it will identify iSNVs that are of high confidence, which can then be used in studies where accurate iSNV determination is of high impact such as in identification of fine scale transmissions.

Interestingly, our results indicated that the frequency of called iSNVs is fairly consistent among different variant callers. This would suggest that identifying true iSNVs is the most challenging part of the process, and once those are identified, the obtained iSNV frequencies are largely reliable. We also notice that when variants within an iSNV position are present at approximately equal frequencies, choice of variant caller might determine which variant is called in the consensus. For instance, in segment 1 position 1868 of sample NA0012484733, the majority variant is T (amino acid L) when LoFreq is used for variant calling, and it is C (amino acid S) when ngs_mapper or iVar are used. This may have a great impact in instances where a consensus sequence is called using the majority rule (only the major variant is reported), as the choice of variant caller alone might determine the nucleotide in that position. This may result in missing the presence of important variants that could have impact on virus fitness, transmission, or resistance to treatment.

Presence of shared iSNVs in different individuals has been suggested to indicate highly accurate transmission links (32). Thus, shared iSNVs have previously been used to aid in determination of transmission chains for several different viruses, including influenza (6, 26–32). However, these studies have likely suffered from challenges of highly accurate iSNV identification. The limitations of consensus sequences alone to resolve epidemiological links, especially in samples collected close in time and space, is very well known and is also evident in all of our phylogenetic trees. Here, the trees are often phylogenetically unresolved, with many nodes having low confidence values. Even in clades with high confidence values, such as in the UMD clade of B/Victoria lineage, the taxa relations within the clade cannot be resolved. Using information from our approach for highly confident iSNV determination, we were able to resolve likely transmission links in our data. It is interesting to note that the viruses from one of the individuals, who belonged to an iSNV-associated transmission (participant 122), displayed identical iSNV positions in samples collected on two consecutive days from this individual. However, the frequencies of these iSNVs had changed drastically over this period of time (Table S2), indicating a very dynamic intra-host viral population, although the impact of sampling bottlenecks cannot be excluded. Consecutive samples (1 day and 2 days apart) from three other individuals were also available. Here, the iSNV positions were not identical over time within a host, and these individuals did not share iSNVs with any other participants. The rate of minor variant turnover within a host may differ between individuals, and it will be a limiting factor for utilization of iSNVs to determine epidemiological links. Even when iSNVs are transmitted between individuals, their rapid turnover may result in loss of signal by the time of sample collection. Thus, shared iSNVs between individuals sampled close in time and space would further point to the specificity of their transmission links.

In summary, we present an approach for removal of amplification, sequencing, and algorithm-related errors for identification of highly confident viral intra-host variants. We apply this approach to a set of influenza samples collected during respiratory surveillance within a cohort of college students in a living learning community, representing close quarter environments with high likelihood of infection transmission chains. Our results point to the prospect and accuracy of utilizing iSNVs for fine-scale tracking of influenza transmission, as well as limitations of this approach. Further studies of intra-host evolutionary dynamics of influenza virus may provide additional insight into the mechanisms of minor variant changes, and aid in construction of improved sampling approaches for fine scale resolution of transmission links.

## MATERIALS AND METHODS

### Samples

This study was approved by the University of Maryland Institutional Review Board and the Human Research Protection Office of the Department of the Army. Electronically signed informed consent was obtained. Informed consent documentation and questionnaire data were collected and stored with REDCap (42). A cohort of college students in a living learning community within a pair of residence halls was recruited for the Prometheus@UMD program sponsored by DARPA in the 2017-18 academic year and monitored prospectively for the occurrence of acute respiratory illnesses (ARIs) (43). Participants who reported ARI-related symptoms were invited to the study clinic to provide mid-turbinate, nasal, and oropharyngeal swab specimens. Swabs were screened for common human respiratory pathogens using a customized TaqMan Array Card^®^ (ThermoFisher, Waltham, MA, USA), which included influenza A and B viruses, adenoviruses, coronaviruses 229E, HKU1, NL63, OC43, paramyxoviruses, respiratory syncytia viruses, and other viral and bacterial targets.

### Plasmids

RNAs were extracted using the RNeasy Mini Kit (Qiagen, Cat. #: 74104) from midturbinate swabs collected in the study that resulted in lowest Ct in the TAC assays for influenza A (Subject ID #239, 2/7/2018) or B (Subject ID #104, 5/1/2018) viruses (Table 1). Using these RNAs as templates, RT-PCRs were carried out with the SuperScript™ III one-step RT-PCR system with Platinum™ *Taq* high fidelity DNA polymerase (ThermoFisher Scientific, Cat. #: 12574-035) and primers and cycling conditions designed to generate full-length cDNA of all influenza virus genome segments (Ref PMID 22528160 and PMID 24501036). The amplicons were gel-purified and cloned into pGEM^®^-T Easy Vectors (Promega, Cat. #: A1360) using standard TA cloning techniques. All plasmid clones were verified by restriction enzyme digestion patterns and Sanger sequencing that covers >500 bp of the insert from both directions.

### Sequencing

Influenza viral RNA was extracted using the QIAamp Viral RNA Mini Kit (Qiagen, USA) in conjunction with Qiagen QIAcube automated system (Qiagen) with QIAamp MinElute Spin kit settings. 200 μl of each clinical specimen was extracted for RNA that was eluted into a final volume of 50 μl. Conventional two steps RT-PCR whole genome amplification were set up in duplicates using 8 pairs of universal primers (44, 45). Plasmids containing clonal (no iSNVs) influenza A and B genome segments from participants in the same cohort were sequenced alongside the influenza samples. Twenty-five μl of viral sample RNA (prepared in duplicates) was used as template for all the 8 reactions and was reverse-transcribed using SuperScript III Reverse Transcriptase (ThermoFisher Scientific, Cat. #: 18080-044). PCR amplification of samples and plasmids was performed using Platinum Taq DNA Polymerase High Fidelity (ThermoFisher Scientific, Cat. #: 11304-102). Amplicons of the same samples were pooled together before purification. AMPure XP beads (Beckman Coulter, Cat. #: A63881) purified amplicons were analyzed for cDNA quality and quantity using TapeStation 4200 (Agilent technologies, Santa Clara, CA) DNA5000 kit (Agilent technologies, Cat. #: 5067-5588, 5589). After TapeStation analysis, ~250 ng of pooled cDNA of each sample was used as input for library preparation using QIAseq FX DNA library kit (QIAgen, Cat.# 180475) or Nextera DNA Flex library Prep kit (illumina, Cat. # 20018705) following the manufactures’ instructions. The purified libraries with different indexes were quantitated using TapeStation D5000 kit and equal molar of each library were pooled together. Pooled libraries were denatured, diluted to an appropriate loading concentration and loaded onto Miseq 600 cycles V3 or Novaseq cluster cartridge of 300 cycle cartridge (illumina, CA, USA) for sequencing.

### Reference mapping and variant calling

Fastqs from Qiagen and Nextera runs for each of the samples’ replicates (S1s and S2s, all 8 segments) and each of the plasmids (P1s and P2s) were merged, and all fastq files were ran on ngs_mapper pipeline for data cleaning through Trimmomatic, and for reference-based genome assembly through BWA-MEM (7, 46). For one of the samples, NA0012484733, three different runs were performed utilizing three different reference genomes for read mapping: B/Louisiana/34/2017 (Victoria lineage, accession numbers: CY249062.1-CY249069.1), B/District Of Columbia/03/2018 (Yamagata lineage, accession numbers: MH584285-MH584292), and a sample self-reference. The self-reference run was conducted through a construction of sample NA0012484733 consensus by initial mapping to the Victoria reference, followed by a re-run with read mapping to the own consensus genome (self-reference). Sample NA0012484733 was the only sample that was investigated using three different references, in order to assess the impact of reference-based iSNV error. All other influenza A and B samples were only ran utilizing self-reference. For influenza A, sample-specific self-reference consensus genomes were created by initial read mapping to the A/Maryland/90/2017(H3N2) genome (accession numbers: CY260950-CY260957). For influenza B, sample-specific self-reference consensus genomes were created by initial read mapping to B/Louisiana/34/2017 (Victoria lineage, accession numbers: CY249062.1-CY249069.1). Three different variant callers were used to call consensus genomes and iSNVs on the data, ngs_mapper’s own variant caller, LoFreq (47), and iVar (48), in order to assess algorithm-related iSNV error. For minor variant (iSNV) calling at a frequency of 1% or higher, a minimum Phred score of 30 and minimum depth of coverage of 1000x throughout the genome were used. The 1% threshold would ensure that each variant was present in a minimum of 10 reads in order for it to be called. iSNVs were determined in four different ways: i) for each of the sample replicates separately (S1, S2), ii) for each of the sample replicates excluding iSNVs found in plasmids, thereby removing systematic sequencing error (S1-P, S2-P), iii) for two sample replicates together by excluding any non-overlapping iSNVs, which would remove PCR and sequencing error (S1+S2), 4) for two sample replicates together while also excluding iSNVs found in the plasmids, thereby removing sequencing, PCR and systematic errors ((S1+S2)-P). All the raw output files were analyzed for presence of iSNVs, giving rise to the results for non-curated data. In addition, all outputs were independently manually curated for a quality check, removing primer-induced mutations, low quality indels, and taking into account sample-specific strand bias where needed, resulting in iSNVs representing the curated data. The non-curated and curated iSNVs were compared to assess the impact of iSNV manual curation-related error. All assembled genomes have been submitted to GenBank under accession numbers listed in Table S3.

### Phylogenetic analyses

We inferred maximum likelihood phylogenies for n = 4 influenza A/H3N2 and n = 11 influenza B full HA-segment sequence data. The HA segment was used to maximize the amount of background reference data and improve phylogenetic inference about distinct introductions of influenza strains into the University of Maryland (UMD) College Park Campus. Background data was obtained by BLAST of study HA segment sequences using the NCBI GenBank database (49). The top 20 BLAST hits per study sequence were selected. We selected additional published influenza A/H3N2 and influenza B full HA segment sequence data sampled 2009-2019 from the NCBI influenza virus database (50). These background data were binned by influenza subtype, collated and de-duplicated before alignment with the UMD study sequences in MAFFT, with manual editing thereafter in MEGA 6.0 (51). This yielded alignments of n = 373 full HA segment influenza A/H3N2 sequences, and n = 762 full HA segment influenza B sequences. Phylogenetic trees for each subtype were inferred by maximum likelihood method in PhyML with nucleotide substitution models selected by JModelTest2 (SYM + G) (52). Root-tip regressions were plotted in TempEst (53), and major regression outliers removed before final phylogenies were inferred in PhyML, with aLRT for node support. All the iSNVs found by the (S1+S2)-P approach, utilizing self-reference for mapping, and confirmed by at least two different algorithms (ngs_mapper’s variant caller and LoFreq), were plotted on the inferred phylogenies for further resolution of sample relatedness.

## DISCLAIMER

Material has been reviewed by the Walter Reed Army Institute of Research. There is no objection to its presentation and/or publication. The View(s) expressed are those of the authors and do not necessarily reflect the official views of the Uniformed Services University of the Health Sciences, Department of Health and Human Services, the National Institutes of Health, the Departments of the Army, Navy or Air Force, the Department of Defense, or the U.S. Government.

## ACKNOWLEDGMENTS

This study was funded by the Military Infectious Disease Research Program, E0002_20_WR and the Armed Forces Health Surveillance Division (AFHSD), Global Emerging Infections Surveillance (GEIS) Branch ProMIS ID: P0116_19_WR and P0140_20_WR. Prometheus-UMD was sponsored by the Defense Advanced Research Projects Agency (DARPA) BTO under the auspices of Col. Matthew Hepburn through agreement N66001-18-2-4015. Material has been reviewed by the Walter Reed Army Institute of Research. There is no objection to its presentation and/or publication. The View(s) expressed are those of the authors and do not necessarily reflect the official views of the Uniformed Services University of the Health Sciences, Henry M. Jackson Foundation for the Advancement of Military Medicine, Inc., Department of Health and Human Services, the National Institutes of Health, the Departments of the Army, Navy or Air Force, the Department of Defense, or the U.S. Government. Dr. Pollett is employed by The Henry M. Jackson Foundation for the Advancement of Military Medicine, Inc. (HJF), a not-for-profit Foundation authorized by Congress to support research at the Uniformed Services University of the Health Sciences (USU) and throughout military medicine. The findings and conclusions in this report are those of the authors and do not necessarily represent the official position or policy of the funding agency and no official endorsement should be inferred. This work was made possible by the collective effort of the Prometheus-UMD investigators listed in Table S4.

## SUPPPLEMENTAL FIGURE LEGENDS

**Figure S1.** Example of workflow overview

**Figure S2.** Number of iSNV positions using the (S1+S2)-P approach per variant caller. A) Influenza A B) Influenza B Victoria. Yellow bar represents positions that were found by all three variant callers. Asterisk notes low viral load samples.

**Figure S3.** Number of iSNV positions in curated sample NA0012484733 when comparing sequencing, reference and algorithm-based errors. A) ngs_mapper variant caller; B) LoFreq variant caller; C) iVar variant caller. Green bars = Self reference; blue bars = Victoria reference; yellow bars = Yamagata reference. S1 = sample replicate 1; S2 = sample replicate 2; S1+S2 = iSNVs confirmed by both S1 and S2; S1-P = sample replicate 1 excluding iSNVs found in plasmids; S2-P = sample replicate 2 excluding iSNVs found in plasmids; (S1+S2)-P = iSNVs confirmed by both S1 and S2 excluding iSNVs found in plasmids.

**Figure S4.** Number of iSNV positions in curated samples. A) influenza A, B) influenza B Victoria CT<30, C) influenza B Victoria CT>30, D) Influenza B Yamagata. Purple bars: ngs_mapper, green bars: LoFreq, orange bars: iVar.

**Figure S5.** Maximum likelihood phylogenetic tree of the influenza B Yamagata HA segment. UMD samples are marked in red text.

