## Supplementary figures and images for "High confidence identification of intra-host single nucleotide variants for person-to-person influenza transmission tracking in congregate settings"

### Supplemental Figures

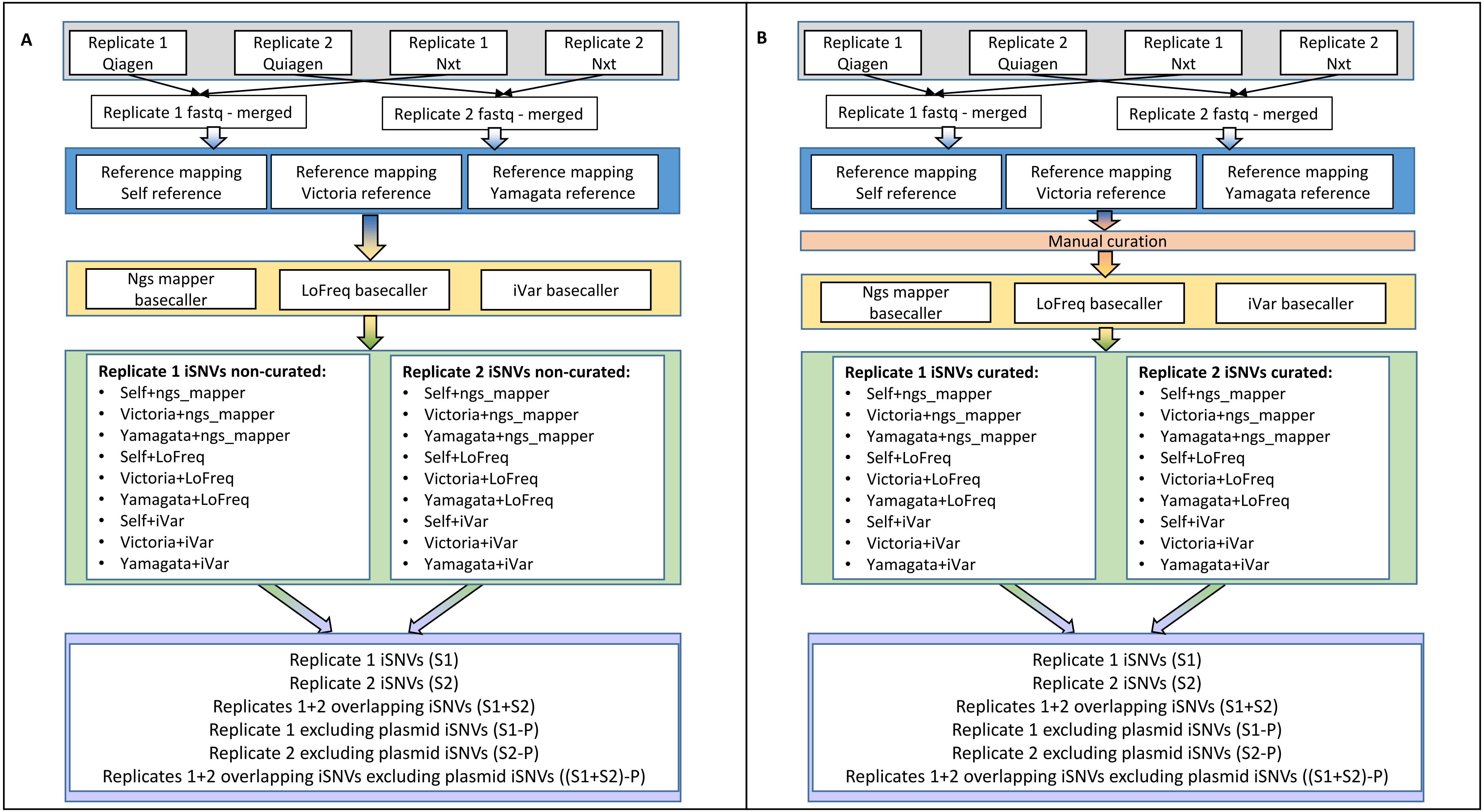

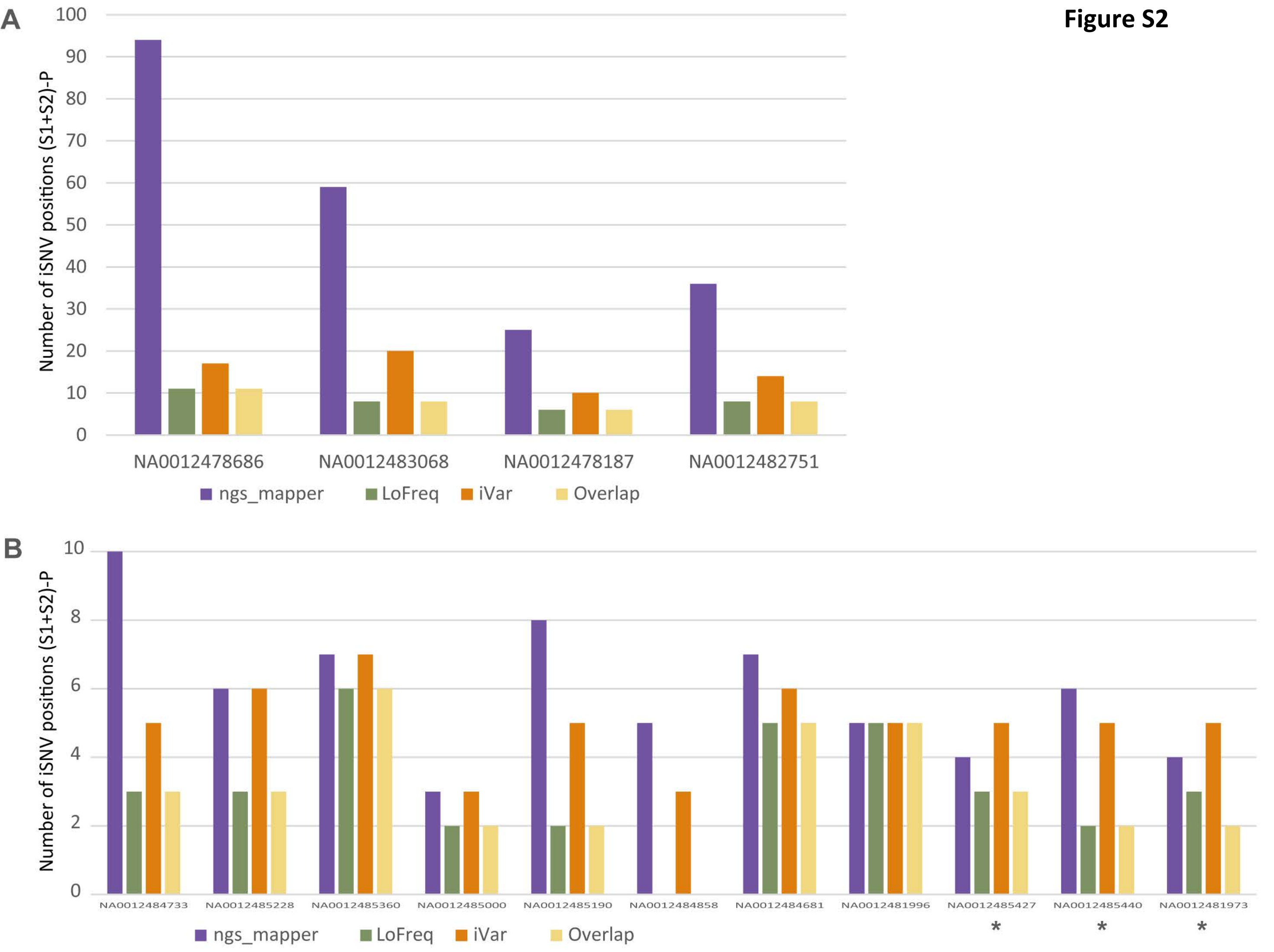

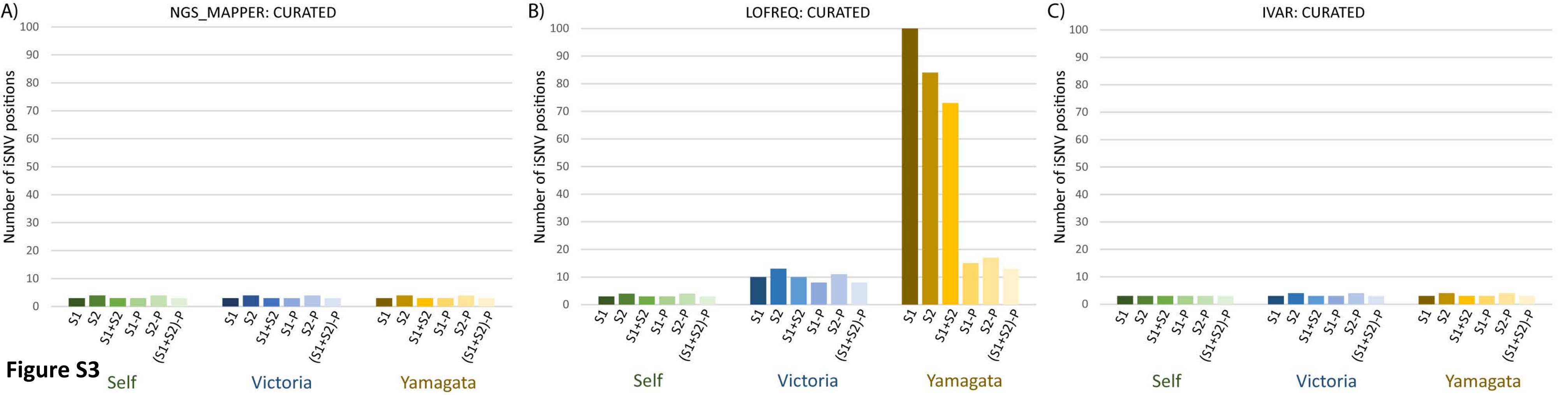

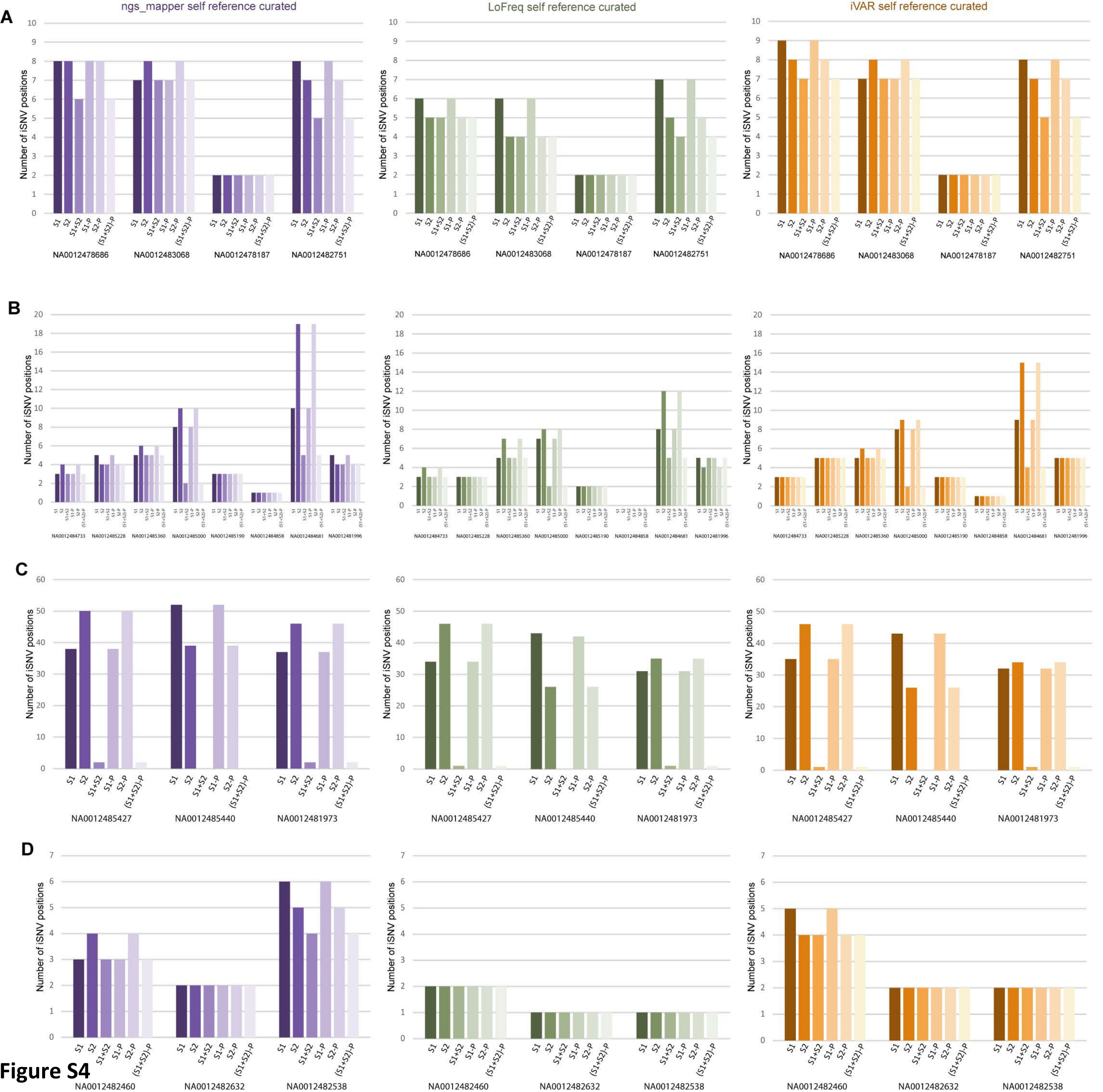

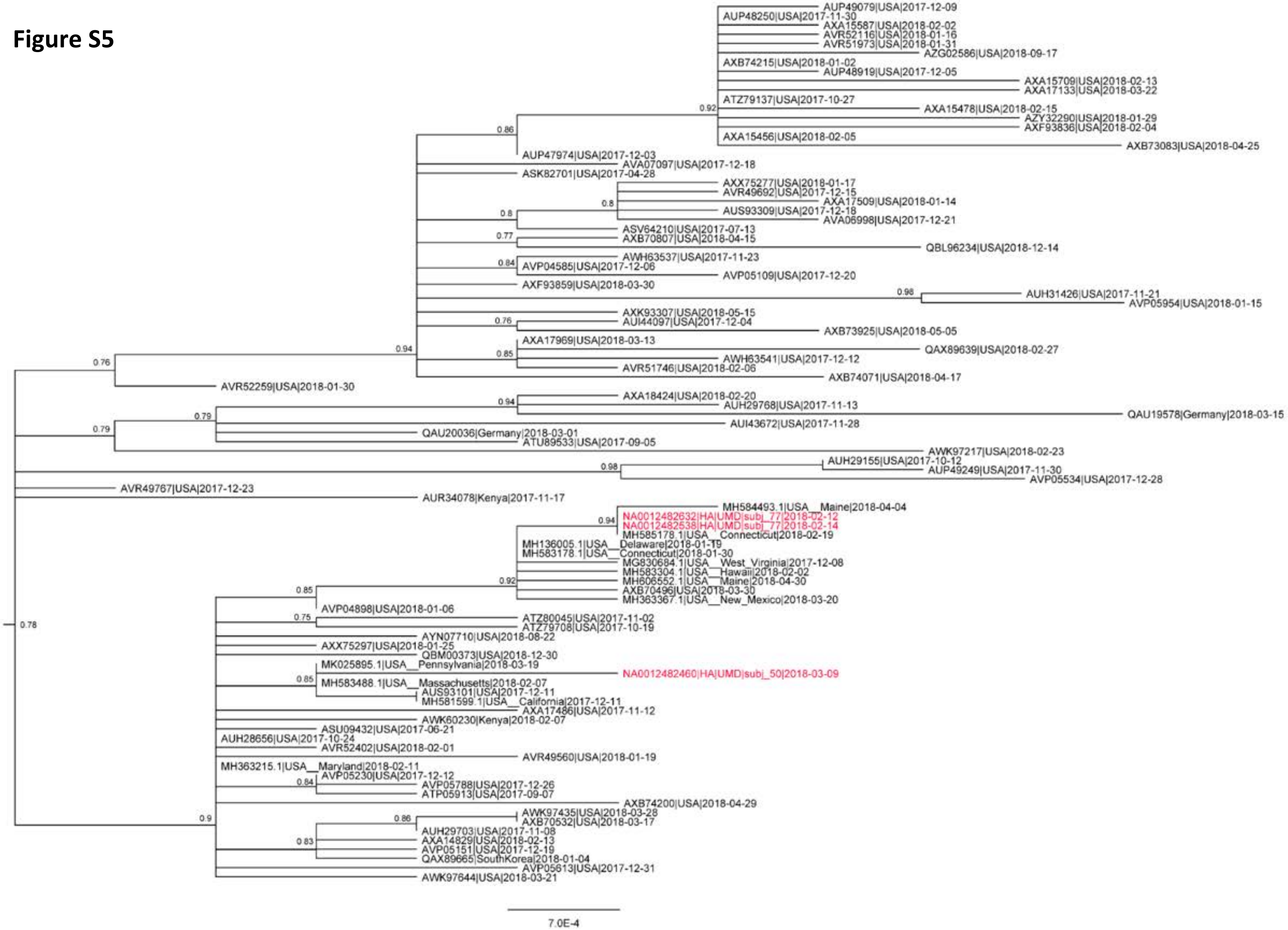
